# Human TSC2 Mutant Cells Exhibit Aberrations in Early Neurodevelopment Accompanied by Changes in the DNA Methylome

**DOI:** 10.1101/2024.06.04.597443

**Authors:** Mary-Bronwen L. Chalkley, Lindsey N. Guerin, Tenhir Iyer, Samantha Mallahan, Sydney Nelson, Mustafa Sahin, Emily Hodges, Kevin C. Ess, Rebecca A. Ihrie

**Author notes:** co-corresponding: (RAI), (KCE).

## Abstract

Tuberous Sclerosis Complex (TSC) is a debilitating developmental disorder characterized by a variety of clinical manifestations. While benign tumors in the heart, lungs, kidney, and brain are all hallmarks of the disease, the most severe symptoms of TSC are often neurological, including seizures, autism, psychiatric disorders, and intellectual disabilities. TSC is caused by loss of function mutations in the *TSC1* or *TSC2* genes and consequent dysregulation of signaling via mechanistic Target of Rapamycin Complex 1 (mTORC1). While TSC neurological phenotypes are well-documented, it is not yet known how early in neural development *TSC1/2*-mutant cells diverge from the typical developmental trajectory. Another outstanding question is the contribution of homozygous-mutant cells to disease phenotypes and whether such phenotypes are also seen in the heterozygous-mutant populations that comprise the vast majority of cells in patients. Using TSC patient-derived isogenic induced pluripotent stem cells (iPSCs) with defined genetic changes, we observed aberrant early neurodevelopment *in vitro*, including misexpression of key proteins associated with lineage commitment and premature electrical activity. These alterations in differentiation were coincident with hundreds of differentially methylated DNA regions, including loci associated with key genes in neurodevelopment. Collectively, these data suggest that mutation or loss of *TSC2* affects gene regulation and expression at earlier timepoints than previously appreciated, with implications for whether and how prenatal treatment should be pursued.

## INTRODUCTION

Tuberous sclerosis complex (TSC) is a genetic developmental disorder characterized by the growth of benign tumors, termed hamartomas, in several organs, as well as neurological symptoms manifesting early after birth. TSC occurs with an approximate frequency of 1 in every 6,000 people worldwide, with wide variation in the severity and incidence of symptoms [1, 2]. 90% of TSC patients are diagnosed with epilepsy, with many patients also exhibiting additional behavioral and neuropsychiatric disorders [3, 4]. The current standard of care for epilepsy includes multiple anti-seizure medications, ketogenic diet, or mTORC1 inhibitors (sirolimus, everolimus), which are generally not curative [5–11]. Select patients with TSC and intractable epilepsy undergo surgical resection of cortical tubers (hamartomas) [12]. Tubers are a classic feature of TSC and are thought to be regions of epileptic activity [13, 14]. Though the gross anatomy of tubers has been described since the 1880s [15], many questions still remain, including the origin and mechanism(s) of formation of these masses. There is a pressing need to understand the cellular origin of tubers, as this could refine the choice of alternative therapeutics for patients and/or identify key windows of opportunity for early treatment.

Dysmorphic neurons and giant cells (also termed balloon cells) are notable diagnostic characteristics of tubers [14] and may provide indications of their etiology. Giant cells have been reported to be simultaneously positive for neural progenitor (nestin and vimentin), neuronal (β- III-tubulin), and glial identity markers (GFAP)[16]. This abnormal multi-lineage co-expression suggests a developmental origin for tubers. Additionally, the stereotypical layers of neurons formed during cortical development are disrupted in tubers, further indicating that early neural development is perturbed [17, 18].

TSC is caused by loss of function mutations in either the *TSC1* or *TSC2* genes, with *TSC2* mutations representing the majority of cases and generally correlating with increased symptom severity [19–23]. Most patients with TSC have *de novo* mutations with no prior family history [21, 24–27]. Of note, about 1/3 of patients inherit a mutant gene in an autosomal dominant pattern. *TSC1* and *TSC2* encode the proteins hamartin and tuberin, respectively [19, 20]. Tuberin is a large protein composed of 1807 amino acids and several functional domains. Hamartin and tuberin form a complex with a recently discovered partner, TBC1D7 [28, 29], and collectively these proteins negatively regulate mammalian target of rapamycin complex 1 (mTORC1)[30, 31]. mTORC1 is a signaling hub known to control protein synthesis and cell growth, among many other cell processes, via its kinase activity affecting downstream targets [32–35]. mTORC1 has also been connected to several additional neurological developmental disorders, which together with TSC are termed mTORopathies [36, 37]. As in TSC, patient brain tissue in other mTORopathies exhibits dysmorphic cells with mixed-lineage phenotypes and aberrant electrophysiology, implying that cell fate specification during development is highly dependent on this signaling pathway [38].

Beyond its direct kinase activity, mTOR has been implicated in the regulation of cellular levels of S-adenosylmethionine (SAM), a universal methyl group donor. While the consequences of SAM level changes on RNA methylation have begun to be explored, it remains unknown how altered mTORC1 activity and SAM levels might affect DNA methylation, a critical modulator of cell fate decisions throughout development. In the developing cortex, mature cell populations are generated sequentially, with stem/progenitor cells first generating a series of neuronal subtypes that populate the layered cortical plate and subsequently generating glia – astrocytes and oligodendrocytes [39–43]. These cell fate decisions are made in part through epigenetic regulation of the genome, with modifications including methylation and chromatin accessibility that mediate transcription of key genes. DNA methylation is catalyzed by DNA methyltransferases (DNMTs) and removed in part by ten-eleven translocation (TET) methylcytosine dioxygenases. The essential role of DNA methylation in cell fate decisions is underscored by DNMT knockout human embryonic pluripotent stem cells (hEPSC), which show either widely dysregulated expression of key genes during development or mass cellular death [44]. During development, DNA methylation is typically associated with gene repression, though recent studies have suggested that DNA methylation does not always directly correspond with gene silencing [45, 46].

Prior human cell-based models of TSC have indicated that the typical pattern and sequence of neurodevelopment is altered in TSC2-mutant cells but have not fully explored the biochemical mechanisms that may drive this altered fate specification [47–50]. Here, we have characterized the earliest stages of cortical development using human iPSCs in both monolayer and organoid culture models of TSC2-deficiency in the brain. TSC2-deficient cultures displayed unexpected co-expression of progenitor cell and neuronal markers along with aberrant increased neuronal activity as early as day 10, a timepoint not addressed in prior studies. This pattern in combination with previous studies showing mTORC1 pathway control of the universal methyl donor suggests global dysregulation of cell fate, prompting a systematic examination of genome regulation through DNA methylation changes.

## RESULTS

### Neural progenitor cell marker protein expression and cytoarchitecture are aberrant in TSC patient-derived cultures

To recapitulate early human neurodevelopment, established isogenic series of TSC2-deficient iPSCs were used. *Patient 77^TSC2^* ^+/LOF^ iPSCs (TSC2 +/LOF) were derived from a TSC patient harboring a disease-causing germline heterozygous C-terminal 6 amino acid deletion in the TSC2 gene (Fig. 1A)[49]. This 6 amino acid deletion occurs in the GTPase activating protein (GAP) domain and results in mutant truncated tuberin protein (Supp. Fig.1A, B). This altered GAP domain is thought to compromise the ability of tuberin to suppress mTORC1 activation by Ras homolog enriched in brain (Rheb) [51]; consequently, mTORC1 activity is elevated in these iPSCs (Sup. Fig.1A, D-E). It is often proposed that “second-hit” somatic mutations occur in a subset of benign TSC tumor cells and can be modeled by homozygous *TSC2* mutation. Thus, in addition to the heterozygous state, experiments were also performed in isogenic lines engineered to have wild type *TSC2* (TSC2 +/+) or homozygous *TSC2* mutation (TSC2 LOF/LOF)(Fig. 1A). A second isogenic pair of lines harboring homozygous wild-type TSC2 (TSC2 WT) or CRISPR engineered knock out of TSC2 (TSC2 KO) were also used to confirm our findings (Fig. 1A)[52].

**Figure 1:**
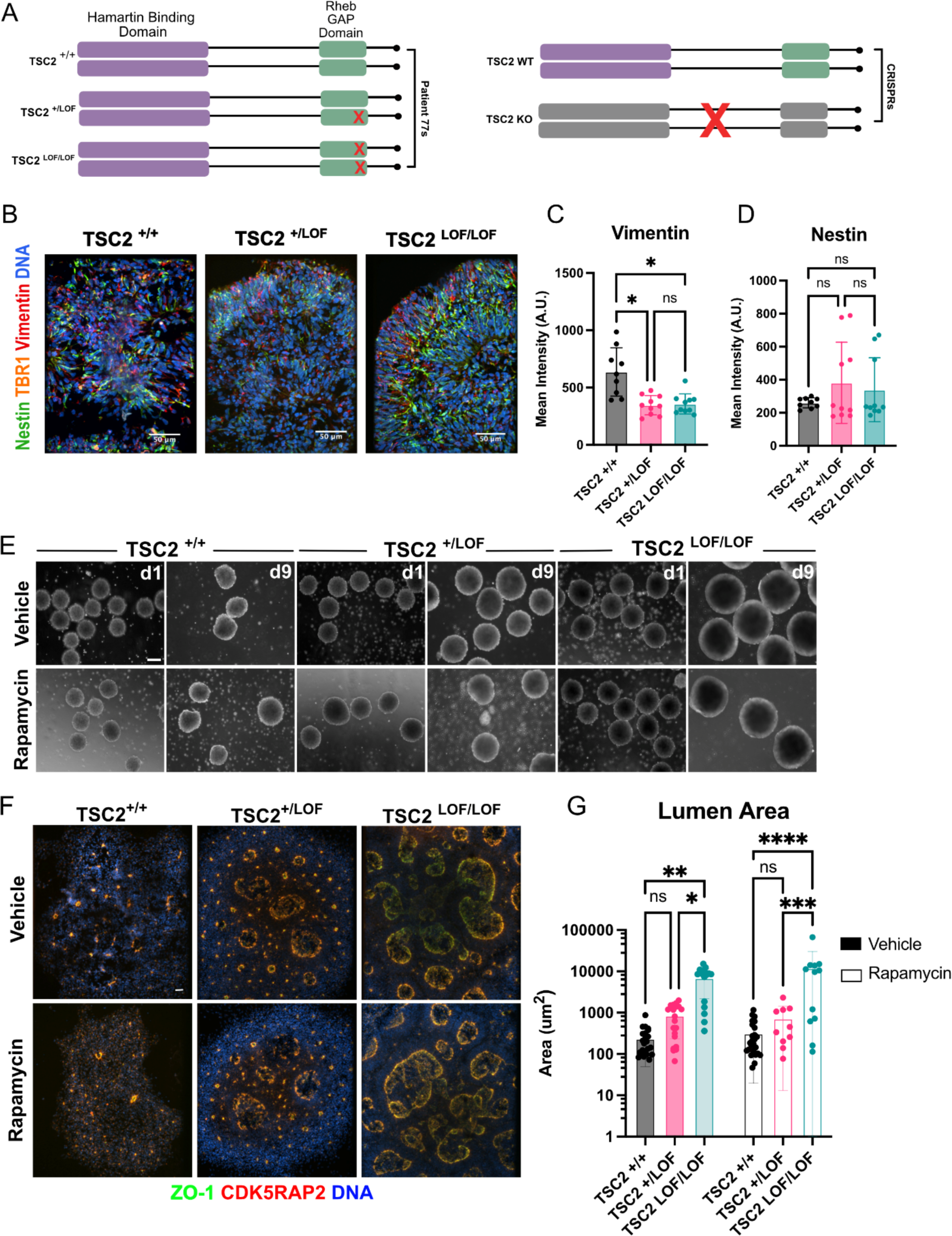
Neural Progenitor marker vimentin is decreased, and neural rosette lumens are enlarged, in TSC2 mutant day 9 organoids. (a) Cartoon depicting the genotypes of the cell lines used in this study. (b) Representative immunofluorescence maximum intensity images of day 9 organoids showing expression of nestin (green), TBR1 (orange), vimentin (red) and Hoechst (blue) to identify neural progenitor cells. Scale bars = 50 µm. (c) Quantification of mean intensity (A.U.) of vimentin. N = 10 organoids across multiple differentiations. +/+ vs. +/LOF * p = 0.0203, +/+ vs. LOF/LOF * p = 0.0229, genotype ** p = 0.0015 [ANOVA, mixed effects]. Error bars = mean with SD. (d) Quantification of mean intensity (A.U.) of nestin. N = 10 organoids across multiple differentiations. No significance [ANOVA, mixed effects]. Error bars = mean with SD. (e) Representative phase DIC microscopy images of day 1 and day 9 organoids treated with vehicle or rapamycin. Scale bar = 200 µm. (f) Representative immunofluorescence maximum intensity images of day 9 organoids treated with vehicle or rapamycin showing expression of ZO-1 (green), CDK5RAP2 (red) and Hoechst (blue) to identify neural rosette lumens. Scale bars = 50 µm. (g) Quantification of area (µm^2^) of day 9 organoids treated with vehicle or rapamycin. N = 25 organoids across multiple differentiations. Vehicle +/+ vs. LOF/LOF ** p = 0.0057, Vehicle +/LOF vs. LOF/LOF * p = 0.0156, Rapamycin +/+ vs LOF/LOF **** p = <0.0001, Rapamycin +/LOF vs LOF/LOF *** p = 0.0002, genotype **** p = <0.0001 [two way ANOVA]. Log 10 scale. Error bars = mean with SD.

These two isogenic series of iPSCs were differentiated for ten days in 3-D organoid culture or 2-D monolayer culture with dual SMAD inhibitors to produce neural progenitor cells (NPC) – the first phase of differentiation for mixed cortical neurons. Neural rosettes and their lumens at this timepoint are thought to model the neural tube *in vitro* [53, 54]. The NPC markers nestin and vimentin were used for an initial confirmation of cell identity. Compared to matched TSC2 +/+ organoids at day 9, patient-derived *TSC2*-mutant NPCs (TSC2 +/LOF and TSC2 LOF/LOF) had significantly decreased intensity of vimentin suggesting either an altered differentiation schedule or lineage trajectory (Fig. 1B, C). There was also a decrease in vimentin intensity in the 2-D monolayer TSC2 mutant cultures (TSC2 LOF/LOF) (Supp. Fig.1F, G).

In concert with changes in vimentin expression, genotype-specific alterations in the size and shape of organoids were evident. While no genotype of organoid was significantly different in overall size measured in diameter at day 1 of culture, TSC2 mutant organoids (TSC2 +/LOF and TSC2 LOF/LOF) had significantly increased in diameter compared to WT (TSC2 +/+) organoids at day 9 of culture (Fig. 1E). Additionally at day 9, TSC2 LOF/LOF organoid lumens had dramatically increased area when compared to TSC2 +/+ lumens. Circularity, a measurement of lumen symmetry, was unchanged in vehicle treated organoids between genotypes. However, circularity was decreased in the rapamycin treated TSC2 mutant (TSC2 +/LOF and LOF/LOF) organoids when compared to TSC2 +/+ lumens showing that TSC2 +/LOF and TSC2 LOF/LOF lumens were not as symmetrical as TSC2 +/+ lumens (Supp. Fig. 1I). All isogenic genotypes showed a decrease in diameter when treated for the entire duration of culture with rapamycin (Fig. 1E). Lumen area did not change significantly in any genotype when the organoids were treated for the entire duration of culture with the mTORC1 inhibitor rapamycin (Fig. 1F, G), suggesting that this feature may be dependent on mTORC1 targets that are less effectively inhibited by rapamycin.

### Neuronal markers and function are prematurely present in TSC2 mutant day 10 neural cultures

The decrease in NPC markers and enlarged lumen morphology of the organoids suggested that the TSC2 LOF/LOF NPCs could be exhibiting either accelerated or aberrant differentiation compared to the TSC2 +/+ NPCs. To examine these possibilities, we next stained for the transcription factor PAX6 and pan-neuronal marker β-III-tubulin. This revealed an increase in the percentage of PAX6-positive cells and also an increase in the percentage of cells expressing β-III-tubulin at this early stage in TSC2 +/LOF and TSC2 LOF/LOF (Fig. 2A-C). This alteration in lineage marker expression was also present in 3D organoids (TSC2 LOF/LOF) and in a second iPSC line with complete loss of tuberin protein (TSC2 KO) (Supp. Fig. 2A-E). Intriguingly, approximately 70% of the β-III-tubulin positive cells found in TSC2 +/LOF and TSC2 LOF/LOF cultures co-expressed PAX6, a decrease from the TSC2 +/+ NPCs which could indicate that the TSC2 mutant NPCs are further differentiated or proceed more rapidly along their developmental trajectory (Fig. 2D, E).

**Figure 2:**
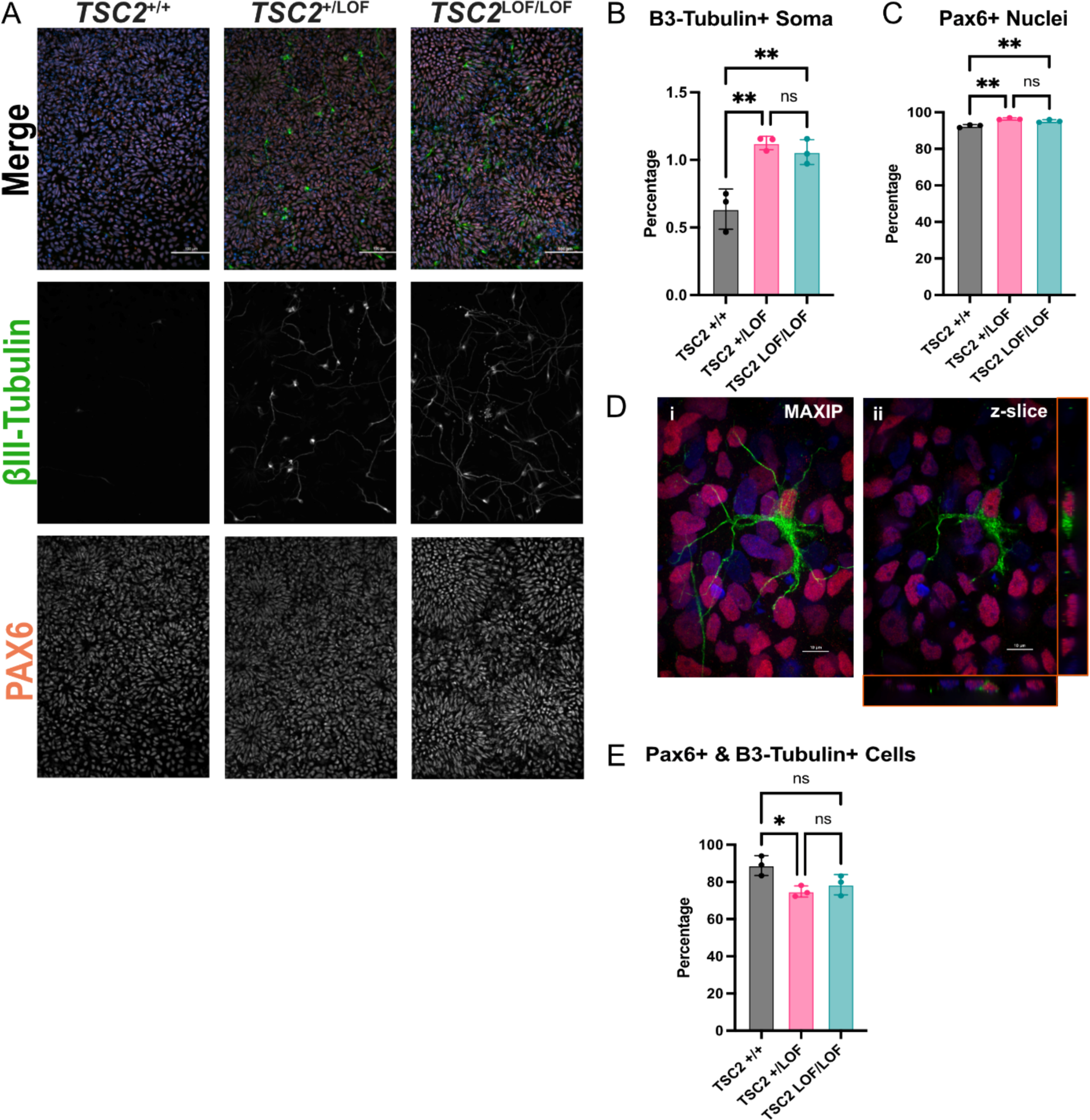
TSC2 homozygous mutants have increased abundance of β-III-tubulin / PAX6 co-positive cells. (a) Representative immunofluorescence maximum intensity images of day 10 monolayer neural cultures showing expression of β-III-tubulin (green), PAX6 (orange), and Hoechst (blue) to identify neural progenitor cells and neurons. β-III-tubulin and PAX6 single channel greyscale images. Scale bars = 100 µm. (b) Quantification of percentage of β-III-tubulin positive soma. N = 3 independent differentiations. Genotype ** p= 0.0025, TSC2 +/+ vs TSC2 +/LOF ** p = 0.0030, TSC2 +/+ vs TSC2 LOF/LOF ** p = 0.0064 [one-way ANOVA with Tukey’s multiple comparisons test]. Error bars = mean with SD. (c) Quantification of percentage of PAX6 positive nuclei. N = 3 independent differentiations. Genotype ** p = 0.0014, TSC2 +/+ vs TSC2 +/LOF ** p = 0.0013, TSC2 +/+ vs TSC2 LOF/LOF ** p = 0.0087 [one-way ANOVA with Tukey’s multiple comparisons test]. Error bars = mean with SD. (d) Representative immunofluorescence images of day 10 monolayer neural cultures showing expression of β-III-tubulin (green), PAX6 (orange), and Hoechst (blue) to showing a cell co-expressing β-III-tubulin and PAX6. i) maximum intensity image. ii) single image of z-slice. Scale bars = 10 µm. (e) Quantification of percentage of β-III-tubulin positive soma that are co-positive with PAX6. N = 3 independent differentiations. Genotype * p = 0.0270, TSC2 +/+ vs TSC2 +/LOF * p = 0.0263 [one way ANOVA with Tukey’s multiple comparisons test]. Error bars = mean with SD.

To explore the function of cultures with increased β-III-tubulin-positive cells, we recorded electrical activity from monolayers using a multi-electrode array machine every other day from day 10 after neural induction to day 28 (Fig. 3A). Neural progenitor cells are not electrically active, and as expected, TSC2 +/+ cultures did not display electrical activity, as measured by number of spikes or mean firing rate, through day 18. However, TSC2 LOF/LOF cultures exhibited low levels of electrical activity throughout the experiment, with activity peaking at day 22 and then decreasing, suggesting a possible cytotoxic effect (Fig. 3B). TSC2 +/LOF cultures had low levels of activity from day 10 to 16, followed by a significant increase of activity from day 16 to day 28, suggesting an increased number of firing neurons in the culture compared to isogenic counterparts (Fig. 3B). Mean firing rate, a second measure of activity, followed a similar pattern as the number of spikes for each genotype (Fig. 3C). Together, this indicates that TSC2 mutant NPCs have accelerated neurogenesis than TSC2 +/+ NPCs.

**Figure 3:**
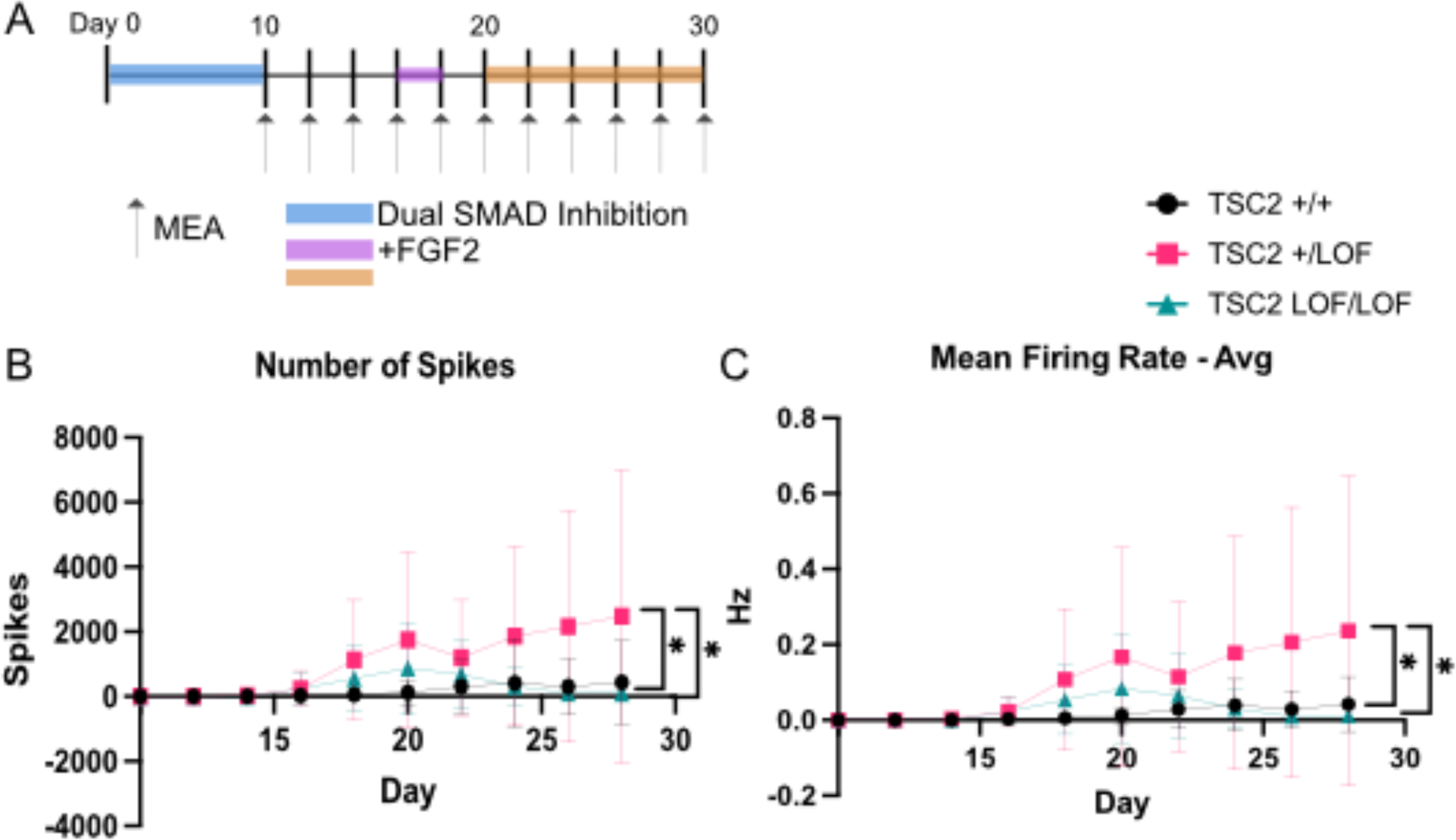
TSC2 mutant neural cells show premature electrical activity. (a) Cartoon showing experimental set up. Arrow represents day MEA recording was done. Color represents media used. (b) Quantification of number of spikes over time. N = 22 wells over 3 independent differentiations. TSC2 +/+ vs TSC2 +/LOF * p = 0.0148, TSC2 +/LOF vs TSC2 LOF/LOF * p = 0.0488, genotype ** p = 0.0083 [one way ANOVA with Tukey’s multiple comparisons test]. Error bars = mean with SD. (c) Quantification of mean firing rate over time. N = 22 wells over 3 independent differentiations. TSC2 +/+ vs TSC2 +/LOF * p = 0.0146, TSC2 +/LOF vs TSC2 LOF/LOF * p = 0.0483, genotype ** p = 0.0081 [one way ANOVA with Tukey’s multiple comparisons test]. Error bars = mean with SD.

Taken together, these results indicate aberrant cell fate decisions in the patient TSC2 mutant cells at an early neural progenitor stage. We hypothesized that the aberrant cell fate decisions we observed are largely the result of epigenetic dysregulation. In particular, dynamic DNA methylation, mediated by DNA methyltransferases and TET enzymes that cause demethylation, is known to be vital in establishing cell identity during development. mTORC1 has been shown to regulate the expression of DNMT3a and DNMT1, as well as the universal methyl donor S-adenosyl-L-methionine. Hence, we stained day 9 organoids for DNMT3a and DNMT1. While we did not observe a significant difference in intensity of DNMT3a in any genotype, we did see a significant decrease in the intensity of DNMT1 protein in TSC2 LOF/LOF (Fig. 4A-C) suggesting a possible change in methylation status in these cells. We examined global methylation in iPSCs and day 10 NPCs for all genotypes. We did not see any changes at the iPSC timepoint, but did observe changes in global methylation in day 10 NPCs for the *TSC2 +/LOF* and *TSC2 LOF/LOF* compared to *TSC2 +/+* (Supp. Fig 4A, B). Consequently, we initiated studies defining the DNA methylome in TSC2 mutant iPSCs and NPCs.

**Figure 4:**
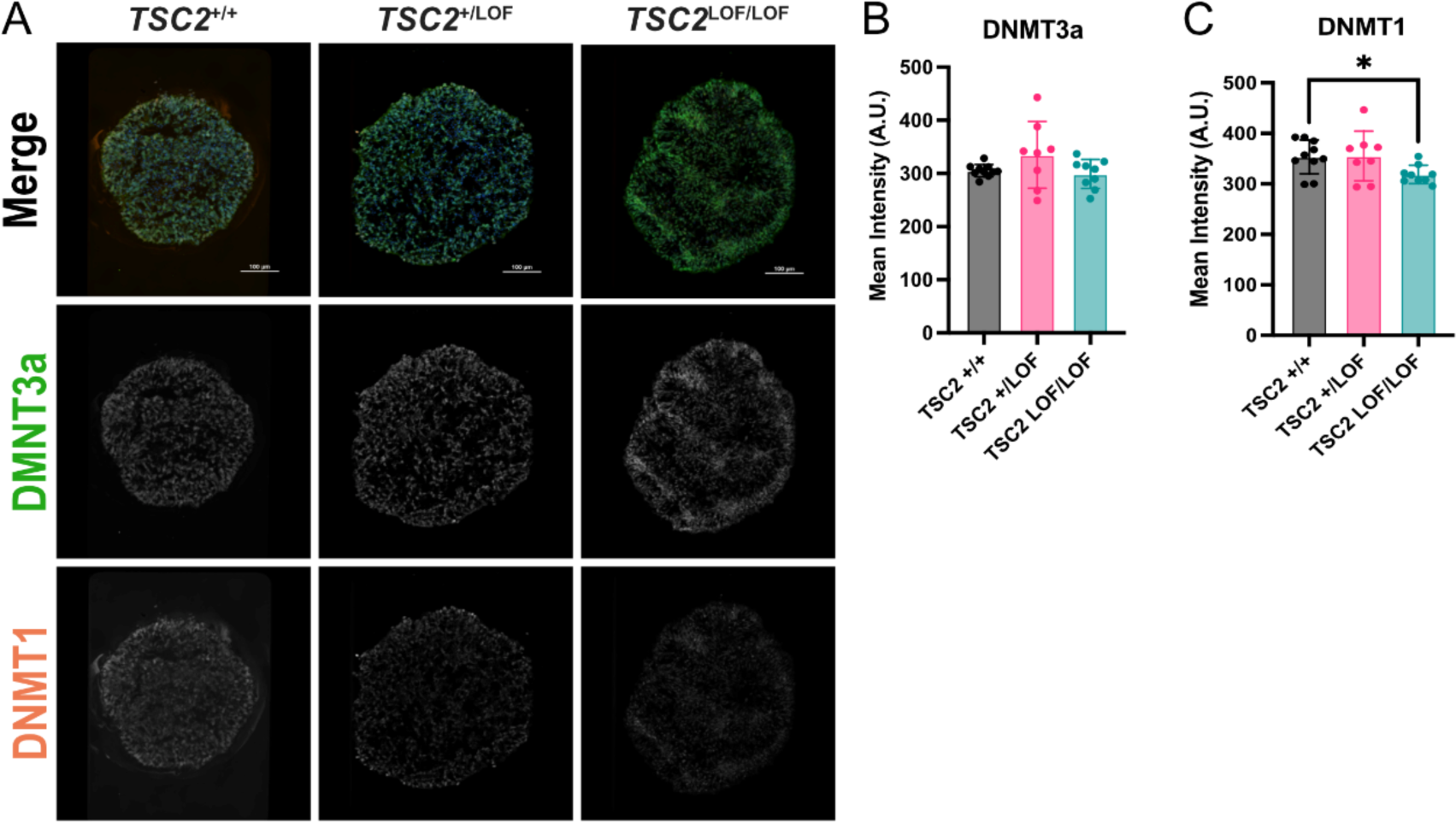
TSC2 mutant day 9 organoids have decreased DNMT1 intensity. (a) Representative immunofluorescence maximum intensity images of day 9 neural organoid cultures showing expression of DNMT3a (green), DNMT1 (orange), and Hoechst (blue). DNMT3a and DNMT1 single channel greyscale images. Scale bars = 100 µm. (b) Quantification of mean intensity (A.U.) of DNMT3a in day 9 organoids. N = 10 organoids across multiple differentiations. No significance [ANOVA, mixed effects]. Error bars = mean with SD. (c) Quantification of mean intensity (A.U.) of DNMT1 in day 9 organoids. N = 10 organoids across multiple differentiations. TSC2 +/+ vs TSC2 LOF/LOF * p = 0.0403 [one way ANOVA, Bartlett’s]. Error bars = mean with SD.

### TSC2 mutant day 10 neural cultures have increased global methylation

We performed whole genome bisulfite sequencing (WGBS) on patient isogenic iPSCs (TSC2 +/+, TSC2 +/LOF, and TSC2 LOF/LOF) and day 10 dual-SMAD neural progenitors derived from these iPSCs (TSC2 +/+, TSC2 +/LOF, and TSC2 LOF/LOF). We first quantified methylation fractions for individual CpG sites genome-wide and defined hypomethylated regions (HMR) using a 2-state hidden Markov model for each sample independently [55]. We identified a range of 41,014-44,614 HMRs per sample. We merged HMRs from each genotype to create a master list of HMRs consisting of both shared and sample-specific HMRs. We examined the distribution of where these HMRs are located within the genome and found that the majority of the HMRs for both iPSCs and day 10 NPCs in all genotypes were within promotor or distal intergenic regions (Supp. Fig. 4C). To identify potential differences in methylation between samples, we averaged methylation levels across all samples for all HMRs. Using unsupervised clustering, we identified 4 groups of HMRs associated with genotype (TSC2 +/+ HMRs and TSC2 LOF/LOF HMRs) or differentiation stage (iPSC HMRs, and NPC HMRs) (Fig.5A, B). Regions in the iPSC and NPC clusters exhibited inverse patterns of methylation levels that were dependent on differentiation timepoint but independent of genotype. In the iPSC HMR cluster, all genotypes at the iPSC stage displayed < 50% average methylation, while those same regions were hypermethylated (HyperMR) in all genotypes at the NPC stage. In the NPC HMRs cluster, regions were hypomethylated in NPC samples in all genotypes, and hypermethylated in all iPSC samples in all genotypes. These differentiation specific differences in methylation were expected between the two timepoints, consistent with previous reports [56]. Separately, regions in the TSC2 +/+ HMRs and TSC2 LOF/LOF HMRs clusters varied by genotype across timepoints, identifying differences in DNA methylation that are primarily associated with *TSC2* genotype.

**Figure 5:**
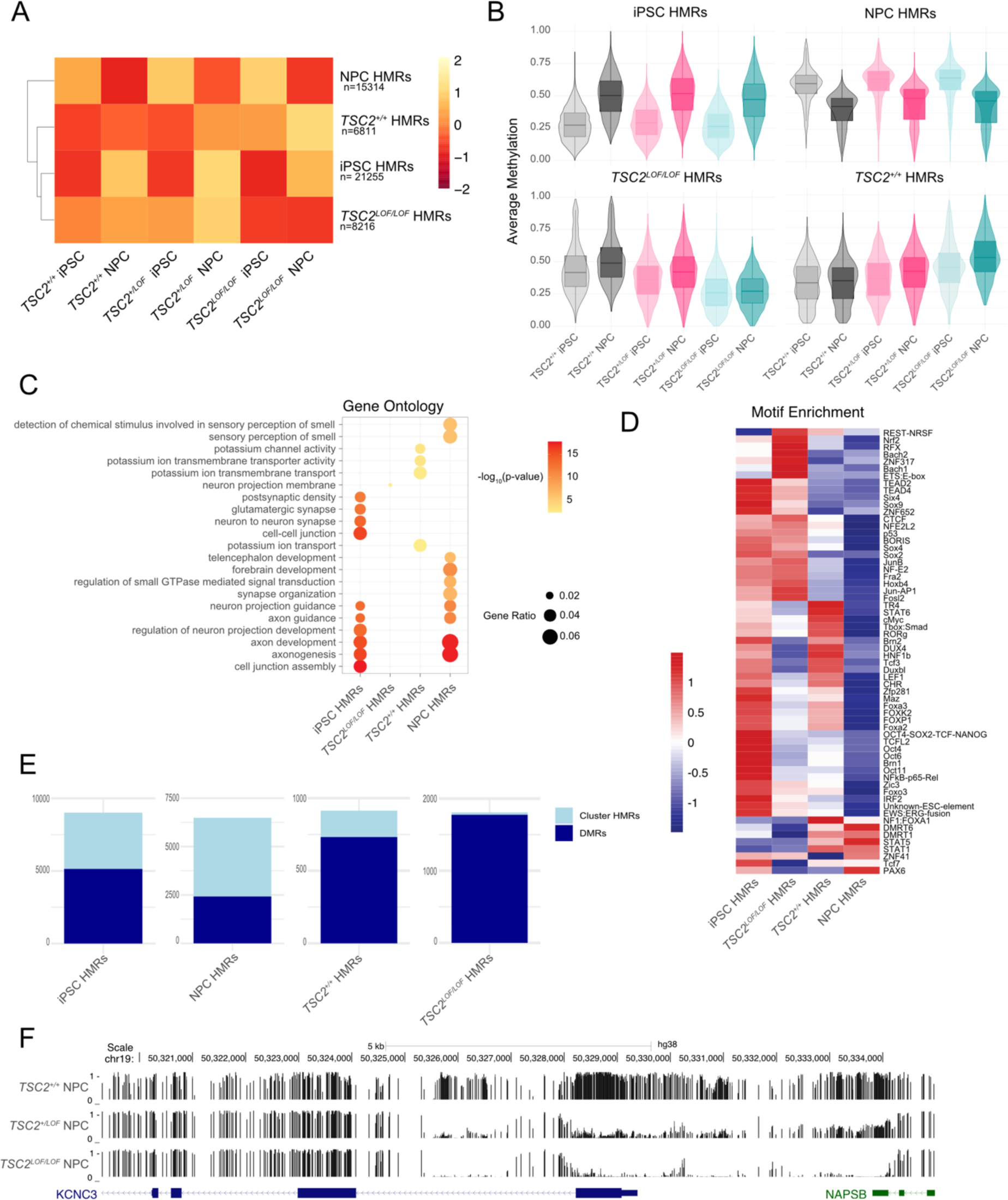
DNA methylation changes with TSC2 genotype. (a) Heatmap of K-means clustering using scale by row feature, input regions were consensus HMR list from all merged samples. Blue indicates hypomethylated and red indicates hypermethylated on scale. (b) Violin plots of average methylation of K-means clustering in a. (c) Gene ontology dot blot from K-means clustering. The n is as follows: iPSCs HMR cluster CC=33932, MF=4110, BP=4003; NPCs HMR cluster CC=1568, MF=1537, BP=1528; TSC2 +/+ HMR cluster CC=0, MF=477, BP=462; TSC2 LOF/LOF HMR cluster CC=816, MF=0, BP=0. (d) Heatmap of motif enrichment for select top most significant motifs from each sample from K-means clustering. Each row is scaled to show the relative enrichment by z-score for the percent fold. (e) Stacked bar plots showing proportion of cluster HMRs that are differentially methylated per cluster. Light blue indicates total HMRs in each cluster, dark blue indicates intersected DMR and HMR. (f) Genomic tracks showing methylation level per CpG with a darker bar indicating higher methylation.

We then examined related gene ontology (GO) terms that distinguished HMRs within each cluster. For the clusters containing TSC2 +/+ HMRs or TSC2 LOF/LOF HMRs, relevant GO terms that emerged were related to potassium ions and potassium channels as well as neuron projection membrane (Fig. 5C). These findings could relate to the aberrant electrical activity found in our TSC2 mutant NPC cultures as well as other studies of TSC2 mutant neurons and patient tissue [49, 50]. We next performed a motif enrichment analysis on each HMR cluster to identify candidate transcription factors that could bind affected regions. Consistent with the clustering analysis, we observed inverse relationships between iPSC HMR and NPC HMR clusters when examining motif enrichment. For example, the pluripotency factor OCT4 is enriched in regions belonging to the iPSC HMR cluster. In contrast, the OCT4 motif is depleted in the NPC HMR cluster (Fig. 5D). We further observed an inverse relationship for motif enrichment for the TSC2 +/+ HMR and TSC2 LOF/LOF HMR clusters again highlighting the differences between the genotypes. This suggests that transcription factor motifs that are hypomethylated in TSC2 +/+ NPCs are depleted in the regions hypomethylated in TSC2 LOF/LOF NPCs, and vice versa. Aberrant methylation of transcription factor binding sites can restrict or permit binding to gene regulatory elements, leading to downstream effects on gene expression and differentiation [57]. Examples of this include motifs for JUN-AP1, JunB, and Fosl2, all factors involved in the maturation of neurons, which have greater enrichment in the TSC2 LOF/LOF HMRs when compared to the TSC2 +/+ HMR regions.

To determine the statistical strength of methylation differences across the HMR clusters, we performed differential methylation analysis on our WGBS data. Differentially methylated regions (DMRs) were calculated by comparing differentiation stage, identifying regions hypomethylated in iPSCs (iPSC lower than NPC) or NPCs (NPC_lower than iPSC). We also compared across genotype to identify regions that were less methylated in TSC2 +/+ samples (WT lower than LOF) or in TSC2 LOF/LOF samples. We determined that 57.12% of the iPSC HMRs and 37.34% of the NPC HMRs contained CpG sites that were differentially methylated (Fig. 5E).

Genotype differences were pronounced, with 80% of the cluster TSC2 +/+ HMRs and all the cluster TSC2 LOF/LOF HMRs containing differentially methylated CpG sites. Several of the DMRs found were in loci for genes relevant in neurodevelopment. An example of one such locus was the gene for potassium voltage-gated channel subfamily C3 (KCNC3) (Fig. 5F). These results indicate that at least part of the observed alterations in cell fate may be driven by altered epigenetic status.

### RNA transcript expression correlate with differentially methylated regions

We then more broadly explored the observed premature neurogenesis phenotype by performing bulk RNA-sequencing of day 10 cultures in each genotype. Principal component analysis showed that samples stratified primarily based on cell line (TSC2 +/+, TSC2 +/LOF, and TSC2 LOF/LOF vs TSC2 WT and TSC2 KO) (Fig. 6A). This would indicate that cell lines with two different *TSC2* mutations should be evaluated separately. RNA-seq analysis showed differential expression of transcripts upregulated in either TSC2 +/+ or TSC2 LOF/LOF. Transcripts upregulated in TSC2 LOF/LOF included *KCNC3*, sodium/potassium proton antiporter (*SLC9A7*), and several zinc finger transcription factors (*ZNF582, ZNF429, ZNF583*) among other genes (Fig 6B).

**Figure 6:**
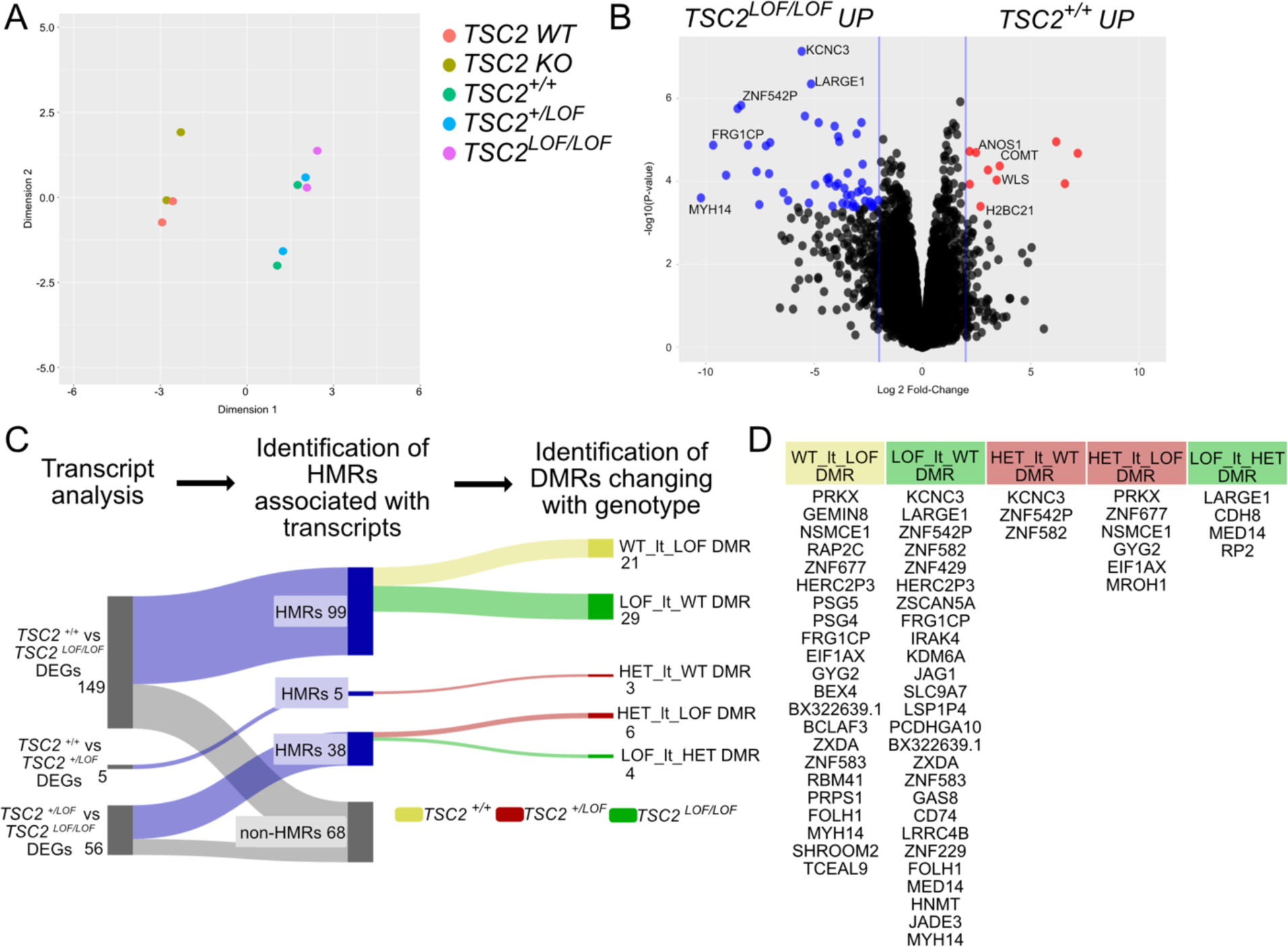
Differentially expressed genes correlate with differentially methylated regions in *TSC2* mutant NPCs. (a) Principal component analysis (PCA) of RNA sequencing day 10 NPC samples. (b) Volcano plot of differentially transcribed genes in *TSC2* +/+ and *TSC2* LOF/LOF. Black dots are genes that are not significant for either fold change or adjusted p-value. Red dots are significantly upregulated in *TSC2* +/+ by both fold change and adjusted p-value. Blue dots are significantly upregulated in *TSC2* LOF/LOF by both fold change and adjusted p-value. (c) Sankey diagram showing correlation of differentially expressed genes (DEGs) to hypomethylated regions (HMR) and differentially methylated regions (DMR). (d) List of differentially expressed genes (DEGs) associated with differentially methylated regions (DMRs).

To determine the potential relationship between differential methylation observed in the WGBS data with differential expression, we quantified expression levels of associated genes via RNA-seq (Fig 6C). We determined whether DEGs were associated with HMRs using the nearest-neighbor approach. All differentially expressed genes (DEGs) found in the RNA-seq data regardless of direction were pooled. 149 DEGs were found when comparing TSC2 +/+ and TSC2 LOF/LOF, 5 DEGs were found when comparing TSC2 +/+ and TSC2 +/LOF, and 56 DEGs were found when comparing TSC2 +/LOF and TSC2 LOF/LOF. We then matched DEGs to annotated HMRs in the WGBS data for day 10 NPCs and further determined associated DMRs for each DEG (Fig 6C, D). While there was not perfect concordance between methylation changes and transcript changes, many DEGs were associated with loci that were differentially methylated across stage and genotype. These DMRs followed the directionality of the RNA-seq data. For example, *KCNC3* transcript was significantly upregulated in TSC2 LOF/LOF compared with TSC2 +/+, and the *KCNC3* locus contained 4 regions that were more hypomethylated in TSC2 LOF/LOF compared with TSC2 +/+, indicating that the change in methylation status for regulatory and coding regions of *KCNC3* resulted in changes in gene expression.

## DISCUSSION

### TSC2 loss of function mutation impacts very early stages of neurodevelopment

TSC is a neurodevelopmental disorder; the debilitating neurological symptoms of TSC, such as autism, psychiatric disorders, and epilepsy, have all been linked to altered number and maturity of neurons, astrocytes, and oligodendrocytes [36, 58]. Though the perinatal impacts of TSC mutations are clinically evident, and later stages of neurodevelopment have been examined using mouse and human-derived models, a gap in knowledge exists regarding whether the earliest stages of neurodevelopment, including neural tube development and patterning, are affected in this disorder. Examining early timepoints in *in vitro* patient iPSC derived neural cultures revealed phenotypic changes in both TSC2 +/LOF and TSC2 LOF/LOF cells. These included alterations in lineage marker expression – accelerated loss of the neural progenitor marker vimentin and an increase in the neuronal marker β-III-tubulin – as well as larger scale cytoarchitectural changes including significantly enlarged lumens in neural rosettes. Collectively, these data indicate disruption of normal neural development in both the heterozygous-mutant context, which represents the majority of cells present in patient tissues, and the homozygous-mutant context, which is variably identified in TSC-associated brain hamartomas [59–61]. Previous data from germline *Tsc2 -/-* mouse models found this genotype to be embryonic lethal at E10.5, with embryos exhibiting unclosed neural tubes [62], and conditional mouse models ablating hamartin (TSC1) or tuberin (TSC2) found an increase in neuron and astrocyte number as well as reduced oligodendrocyte number and decreased myelination [63–66]. The premature appearance of astrocytes has also been reported in human iPSC-derived models of TSC at timepoints later than those examined here [47, 48, 67]. Our findings are consistent with these reports but indicate a more pervasive and early effect of *TSC2* mutation on neural lineage commitment than previously appreciated. These data argue that a closer examination of very early timepoints during *in vitro* differentiation is needed in models of TSC as well as other mTORopathies, expanding on existing studies that have focused extensively on cell phenotypes reflecting mid-gestation stages. A question raised by these human cell model studies is what timepoints may be amenable to clinical intervention, either through early genetic testing or other approaches.

Further research is required to distinguish competing mechanistic possibilities underlying the phenotypes observed here. iPSC-derived models of a wide variety of neurodevelopmental disorders, both mTORopathies and others, have reported enlarged or dysmorphic lumens in neural rosette structures. Cross-species studies probing genetic changes underlying differing cortical size and gyrification have also highlighted both mTOR activity and regulators of cell shape as processes that are modulated as brain size and organization changes. However, the directionality of this relationship is not clear. Though enlarged lumens can be found in several models of neurodevelopmental disorders, it is not clear if alterations in lumen size and cellularity are a cause or consequence of altered neural fates [68]. Abnormal morphology has been connected to altered cell fate decisions in neural rosettes [69], and it is possible that the morphological changes seen in the TSC2 +/LOF and TSC2 LOF/LOF organoids might be directly impacting cell fate decisions. Alternatively, TSC-mutant cells may undergo accelerated progression through differentiation programs resulting in altered cytoarchitecture and large-scale disorganization of rosettes. Given possible mechanistic links between mTOR and transcriptional/translational programs affecting differentiation or cytoskeletal organization, it is particularly interesting that treatment with rapamycin did not ameliorate the observed cytoarchitectural phenotypes. Multiple studies have shown that rapamycin is effective at decreasing phosphorylation of S6, but largely ineffective at durably inhibiting phosphorylation of other mTORC1 downstream targets including 4E-BP1/2. This suggests that the changes in protein expression and morphology are not an S6 driven process, despite the widespread usage of p-S6 as a proxy for mTOR activity in studies of brain development.

### TSC2 loss of function mutation alters the epigenome in early neural development

Epigenetic regulation is vital for proper development. In particular, DNA methylation is important for cell fate decisions of cortical neurons, with hypermethylation serving as a key mechanism for restricting transcription factor access to gene loci as differentiation proceeds [70]. We have identified differences in DNA methylation across *TSC2* genotypes during early neuronal differentiation. Wild-type day 10 NPCs exhibited hypermethylation across many regions that were hypomethylated in matched iPSCs, consistent with expected developmental patterns, and many of these regions were enriched for motifs of canonical pluripotency or neural differentiation factors, including OCT4 and SOX2. In day 10 NPCs, several regions that were hypermethylated in wildtype day 10 NPCs were instead hypomethylated in *TSC2*-mutant cells, suggesting the potential for aberrant transcription in the mutant cells. This difference in methylation status could underlie some of the aberrations in neural development noted in the TSC2 mutant organoids and monolayer cultures, as in the case of KCNC3 which expression changes could reflect in electrical changes examined. We noted that the strongest pattern of differential methylation between genotypes was in the day 10 NPCs compared to the iPSCs, indicating that the effect of the mutation becomes more pronounced as development proceeds. These methylation changes could also underlie some of the aberrations in protein expression seen in this study with phenotypical changes occurring at day 10 of differentiation. Congruent with this idea, of differentially expressed genes that were associated with differentially methylated regions, all the differentially methylated regions showed changes with a directionality consistent with the observed transcript level changes, indicating that changes in DNA methylation have widespread effects on transcription in TSC mutants.

Interestingly, RNA sequencing of day 10 NPC cultures did not show differential transcription for the proteins that were differentially expressed. mRNA expression for *VIM* (vimentin), *NES* (nestin), *PAX6*, and *TUBB3* (β-III-tubulin) did not change significantly between any genotypes. This could be due to the relatively small proportion of cells with altered expression of these proteins, which would not be well represented in bulk sequencing inputs. A second possibility is that some of protein expression changes seen in this study are not a product of transcriptional changes but rather post transcriptional regulation including preferential translation, protein modification, or protein degradation. Preferential translation could be a possible mechanism driving these phenotypes; mTORC1 is canonically involved in the phosphorylation of targets that enable cap-dependent translation, including 4E-BP1/2, and preferential translation has been implicated in the process of activation and differentiation of neural stem cells in the adult brain [71]. Additionally, approximately half of the differentially expressed genes found in RNA sequencing were not clearly associated with differentially methylated regions, indicating additional layers of gene regulation, potentially including elevated transcription driven by the transcription factors found above. For example, RNA-sequencing showed upregulation of several zinc finger transcription factors in the TSC2 LOF/LOF and TSC2 KO which could also result in changes of cell fate decisions by aberrantly turning genes on or off.

The methyl groups used in epigenetic regulation originate from S-adenosyl methionine (SAM), which serves as the major donor for nucleic acid and protein substrates. Recently, mTORC1 has been found to regulate this universal methyl donor to increase the methylation of m^6^A RNA, which stimulates protein synthesis. Additionally, mTORC1 was also found to regulate the genome wide histone H3 trimethylation at lysine 27 (H3K27me3). However, DNA methylation in cells with elevated mTORC1 activity had not previously been explored. The findings here indicate widespread alterations during early stages of neural fate commitment, including altered expression of zinc finger transcription factors and proteins important for neuronal function, notably KCNC3 and SLC9A7. Future studies expanding beyond the DNA methylome to additional epigenetic marks, such as histone methylation and acetylation, will be necessary to provide a more comprehensive picture of the epigenetics underlying aberrant neurodevelopment in TSC. Nonetheless, this study, which identifies differences in DNA methylation in both heterozygous and homozygous mutant cells, is an important step to illuminate mechanisms behind abnormal brain development with aberrant neural cell commitment. Such knowledge will help define windows of opportunity for future therapeutic approaches for patients with TSC.

## METHODS

### Resource Availability

#### Corresponding Author

Further information and requests for resources and reagents should be directed to and will be fulfilled by the co-corresponding authors, Rebecca A. Ihrie and Kevin C. Ess.

#### Materials availability

This study did not generate new unique reagents.

Data sets can be accessed at GSE264590 and GSE269086.

A stepwise description of code and tools used in this study can be found at github.com/ihrie-lab/TSC2-DNAme

### hiPSC Cell Culture

iPSCs were grown as colonies on Matrigel (Corning) coated 6 well plates or glass bottom 35mm plates (Cellvis) in mTeSR1 medium (StemCell Tech), replaced daily, maintained at 37°C and 5% CO_2_, and passaged as needed with ReLSR (StemCell Tech).

### Mixed Cortical Neuron Differentiation

Neuronal cultures were differentiated and cultured as previously described [72, 73]. 3-D cultures were formed by seeding 3 × 10^6^ hiPSCs into an Aggrewell800 (StemCellTech) well. Cells were allowed to form spheres for 24 hours and then transferred to 6-well plates to grow in suspension. Suspension spheres were agitated continuously on a rotating platform at 80 rpm.

Media formulations followed exactly as described above [73] in order to be able to compare timepoints between 2-D and 3-D cultures.

### Whole Cell Extract

Cells were washed with 1X PBS with 10 µM sodium vanadate. Cells were lysed using lysis buffer (1% Triton X-100 in STE [100mM NaCl, 1mM EDTA, 10mM Tris pH 8.0], PhosSTOP (Sigma Aldrich), PIC (Sigma Aldrich), 50µM MG132 (Sigma Aldrich), 1µM PMSF (Sigma Aldrich)) added directly to the plate followed by scraping of the cells. Lysate was then sonicated for 3 seconds on power 3 at 4°C. Lysate was centrifuged for 15 minutes at 16,000 x g at 4°C. Supernatant was kept at -80°C until further analysis.

### Immunoblotting

Samples were prepared by mixing whole cell extracts with 4X SDS (Bio-Rad, Hercules, CA) and then boiled for 5 minutes at 105°C. Gel electrophoresis was performed on a 4-12% Bis-Tris gel (ThermoFisher) with 1X NuPage MOPS running buffer (ThermoFisher) at a constant voltage of 140V for 90 minutes. Proteins were transferred to a PVDF membrane in transfer buffer on ice in the cold room (4°C) overnight at a constant current of 33 mA. Post-transfer, the membrane was stained with Ponceau S solution (Sigma Aldrich) for 5 minutes to determine total protein loaded into gel lanes. Membranes were washed 3X in ddH2O. Membranes were blocked in 5% non-fat milk (RPI Corp, Mount Prospect, IL) in Tris-buffered saline (Corning) with 0.1% Tween 20 (Sigma Aldrich)(TBS-t) for 1 hour at room temperature with agitation. All primary and secondary antibodies were diluted in TBS-t with 5% non-fat milk, and the membrane was washed 3X with 0.1% TBS-t afterwards. Antibodies are listed in the table 1. Primary antibodies were incubated overnight at 4°C with agitation. Secondary antibodies were incubated for 1 hour at room temperature with agitation. The membrane was imaged using ECL developing reagents (ThermoScientific) and CCD Imager – AI600 (General Electric).

**Table 1:** Antibodies used.

| Antibody | Manufacturer | Catalog | Dilution | Application |
| --- | --- | --- | --- | --- |
| p-4E-BP1/2 (Thr 37/46) | Cell Signaling Technologies | 2855 | 1:1000 | Western Blot |
| 4E-BP1 | Cell Signaling Technologies | 9644 | 1:1000 | Western Blot |
| CDK5RAP2 | Invitrogen | 702394 | 1:500 | Immunostaining |
| β-III-Tubulin | Cell Signaling Technologies | 4466 | 1:300 | Immunostaining |
| DNMT1 | Cell Signaling Technologies | 5032 | 1:100 | Immunostaining |
| DNMT3a | Novus | NB120-13888 | 1:500 | Immunostaining |
| Hamartin | Cell Signaling Technologies | 6935 | 1:1000 | Western Blot |
| Nestin | Stem Cell Technologies | 60091.1 | 1:500 | Immunostaining |
| PAX6 | Cell Signaling Technologies | 60433 | 1:200 | Immunostaining |
| TBR1 | Cell Signaling Technologies | 49661 | 1:400 | Immunostaining |
| TUBERIN (TSC2) | Cell Signaling Technologies | 4308 | 1:2000 | Western Blot |
| p-S6 Ribosomal Protein (Ser 240/244) | Cell Signaling Technologies | 5364 | 1:1000 | Western Blot |
| S6 Ribosomal Protein | Cell Signaling Technologies | 2217 | 1:1000 | Western Blot |
| Vimentin | Millipore | AB5733 | 1:1000 | Immunostaining |
| ZO-1 | Invitrogen | 33-9100 | 1:500 | Immunostaining |
| Donkey anti-Mouse IgG, Alexa Fluor 488 | Invitrogen | A21202 | 1:1000 | Immunostaining |
| Goat anti-Rabbit IgG, Alexa Fluor 568 | Invitrogen | A11036 | 1:1000 | Immunostaining |
| Anti-Chicken IgG, Alexa Fluor 568 | Invitrogen | A11041 | 1:1000 | Immunostaining |
| Anti-Chicken IgG, Alexa Fluor 647 | Invitrogen | A21449 | 1:1000 | Immunostaining |
| Goat anti-mouse IgG, HRP | Invitrogen | 62-6520 | 1:5000 | Western blot |
| Goat anti-rabbit IgG, HRP | Invitrogen | 65-6120 | 1:5000 | Western Blot |

### Immunofluorescence

Cells were fixed by incubation in either 4% Paraformaldehyde for 10 minutes at 4°C or 100% methanol for 10 minutes at -20°C. Fixed samples were blocked with blocking buffer [PBS(Corning), 1% Normal Donkey Serum (ThermoScientific), 1% BSA (Sigma Aldrich), 0.1% Triton X-100] for 1 hour at room temperature. Primary antibodies were diluted in blocking buffer and then incubated overnight at 4°C. Secondary antibodies were diluted in blocking buffer and then incubated 1 hour at room temperature in the dark. Antibodies are listed in the table 1. Images were acquired using a Prime 95B camera mounted on a Nikon spinning disk microscope using a Plan Apo Lambda 20x objective lens. The software used for image acquisition and reconstruction were NIS-Elements Viewer (Nikon) and ImageJ (FIJI).

### Image Analysis

The majority of images were analyzed using NIS-Elements (Nikon). Quantification of β-III-tubulin+ soma was performed using a custom python script. First, a maximum intensity projection was performed on acquired z-stack images. The Hoechst channel was used to segment the nuclei of all cells using a custom Stardist model trained on manually annotated training and testing images randomly selected from a dataset of the same cell type and growth conditions. These labels were used in conjunction with the appropriate channel images to quantify the expression of markers that exhibited nuclear localization. Next, a series of manipulations were performed on the β-III-tubulin image in order to threshold only the β-III-tubulin+ soma and ignore β-III-tubulin+ processes overlapping β-III-tubulin-cells. The β-III-tubulin image was initially thresholded for all putative β-III-tubulin+ pixels. The resulting binary mask was eroded by 8 pixels and dilated by 8 pixels in series. The result of the erosion and dilation steps is the loss of masked regions with less than 17 pixels of width. Practically, these steps allowed for the removal of the relatively thin β-III-tubulin+ processes. Next, a simple segmentation script labeled each β-III-tubulin+ soma and recorded their centroid coordinates. Finally, the coordinates for each labeled β-III-tubulin+ soma were matched to the nearest labeled Hoechst+ cell. This matching allowed for a single-cell quantification of β-III-tubulin soma signal along with the other markers of interest. Two-way ANOVA and student t-tests were run using Prism (GraphPad) on the collected data.

### Multi-Electrode Array

Induced pluripotent stem cells (iPSCs) were differentiated into neural progenitor cells (NPCs) following protocol previously described (Snow et al. 2020). One day prior to cell plating, MEA wells were coated with a 0.1% polyethyleneimine solution to promote cell adherence to the electrode recording active zone. On day 8 of neural differentiation, NPCs were plated to a 48-well Axion CytoView MEA plate at a seeding density of 100,000 cells/well using a droplet method to ensure that all cells were attached to the active zone. 8 MEA wells were plated per genotype. Subsequently, each well was given 300 uL of NIM supplemented with laminin (10 ug/mL). 3 mL sterile water was applied to each humidity chamber adjacent to recording wells to ensure that cells did not dry out. On day 9, NPCs were fed with BrainPhys Complete Maturation Medium [BrainPhys basal medium (Stem Cell Technologies), 1X B-27+ Neuronal Culture System (50X) (Gibco), 1X N-2 Supplement (100X) (Gibco), 20 ng/mL BDNF (Peprotech), 20 ng/mL GDNF (Peprotech), 1 mM dibutyryl-cAMP (Sigma), and 200 nM Ascorbic Acid (Fisher)]. From day 10 to day 28, cells were half-fed every 2-3 days with BrainPhys Complete Maturation Medium.

Beginning on day 10 until day 28, electrophysiological activity of NPCs was recorded using an Axion Maestro Pro. Prior to recording, the recording plate was left to acclimate for 10 minutes in the MEA chamber at 37°C and 5% CO2. Neural activity was recorded for 10 minutes every other day. Recordings of spontaneous electrical activity from each well were conducted using the Neural Module on the Axion AxIS Navigator software. On days where plate recording and cell feeding coincided, NPCs were fed after recordings were completed to avoid potential nutrient influences on electrical activity of NPCs. All recording files were batch processed to assess the rate of spontaneous neural firing and network bursting across wells for each genotype.

### Whole Genome Bisulfite Sequencing (WGBS)

WGBS was performed as previously described in [74]. Briefly, genomic DNA was extracted using phenol-chloroform extraction from 8-12 million cells. Purity and integrity of genomic DNA was confirmed using Nanodrop and Tapestation (Agilent). 50ng of genomic DNA was then incubated with Tn5 assembled with adaptor oligos(produced in-house) and reaction buffer (5x Tris-DMF) at 55C for 8 minutes. Resulting fragments were purified prior to gap repair and oligo replacement. Bisulfite conversion of repaired DNA fragments was performed using the Zymo-Gold Methylation Kit according to the manufacturer’s instructions. Libraries were amplified and purified prior to sequencing on a NovaSeq.

### WGBS analysis

Raw sequencing reads were first trimmed using Trim-Galore. Trimmed reads were then mapped on to hg38A genome using Abismal [75, 76]. Mapped reads were then processed using MethPipe suite of tools [77]. Methylation levels were calculated at symmetrical CpGs using the `methcounts` command. CpGs with a read count less than five were removed for quality control. HMRs from all samples were concatenated and merged to generate a consensus region list. Average regional methylation at each region was quantified using the `roimethstat` command in MethPipe. Resulting values across all consensus regions were clustered using K-means clustering and named after dominating characteristics of the cluster. Gene ontology was performed using ChipSeeker [78, 79] and motif enrchment analysis was performed using HOMER [80]. Differentially methylated regions (DMR) were calculated using the `methdiff` command in the MethPipe suite of tools.

### RNA extraction

NPCs were collected and pelleted at day 10 of differentiation. Pellets were stored at -80°C until all samples had been collected. All samples of RNA were extracted at the same time. RNA was extracted following the protocol in the RNeasy Mini Kit (Qiagen). RNA extraction included on-column treatment with DNase I following the optional portion of the RNeasy kit using the DNase I kit (Qiagen).

### RNA Sequencing and Analysis

Samples were prepared according to library kit manufacturer’s protocol, indexed, pooled, and sequenced on an Illumina NovaSeq X Plus. Basecalls and demultiplexing were performed with Illumina’s DRAGEN and BCLconvert version 4.2.4 software. RNA-seq reads were then aligned to the Ensembl release 101 primary assembly with STAR version 2.7.9a. Gene counts were derived from the number of uniquely aligned unambiguous reads by Subread:featureCount version 2.0.3. Isoform expression of known Ensembl transcripts were quantified with Salmon version 1.5.2. Sequencing performance was assessed for the total number of aligned reads, total number of uniquely aligned reads, and features detected. The ribosomal fraction, known junction saturation, and read distribution over known gene models were quantified with RSeQC version 4.0.

All gene counts were then imported into the R/Bioconductor package EdgeR and TMM normalization size factors were calculated to adjust for samples for differences in library size. Ribosomal genes and genes not expressed in the smallest group size minus one samples greater than one count-per-million were excluded from further analysis. The TMM size factors and the matrix of counts were then imported into the R/Bioconductor package Limma. Weighted likelihoods based on the observed mean-variance relationship of every gene and sample were then calculated for all samples and the count matrix was transformed to moderated log 2 counts-per-million with Limma’s voomWithQualityWeights. The performance of all genes was assessed with plots of the residual standard deviation of every gene to their average log-count with a robustly fitted trend line of the residuals. Differential expression analysis was then performed to analyze for differences between conditions and the results were filtered for only those genes with Benjamini-Hochberg false-discovery rate adjusted p-values less than or equal to 0.05.

## Acknowledgements

We would like to thank members of the Ess and Ihrie labs for their insightful feedback.

We would like to thank the Vanderbilt Cell Imaging Shared Resource and Nikon Center of Excellence. Microscopy experiments/data analysis/presentation were performed in part through the use of the Vanderbilt Cell Imaging Shared Resource.

We thank the Genome Technology Access Center at the McDonnell Genome Institute at Washington University School of Medicine for help with genomic analysis. The Center is partially supported by NCI Cancer Center Support Grant #P30 CA91842 to the Siteman Cancer Center from the National Center for Research Resources (NCRR), a component of the National Institutes of Health (NIH), and NIH Roadmap for Medical Research. This publication is solely the responsibility of the authors and does not necessarily represent the official view of NCRR or NIH.

## Contributions

MBLC – data acquisition, data analysis, data visualization, experimental design, conceptualization, writing – original draft, writing – review and editing

LNG - data acquisition, data analysis, data visualization

TI - data acquisition, data analysis

SM - data acquisition, data analysis, data visualization

SN - data acquisition

MS - resources EH – resources

KCE – experimental design, conceptualization, resources, funding acquisition, writing – review and editing

RAI – experimental design, conceptualization, resources, funding acquisition, writing – review and editing

