## Supplemental Figures for "Human TSC2 Mutant Cells Exhibit Aberrations in Early Neurodevelopment Accompanied by Changes in the DNA Methylome"

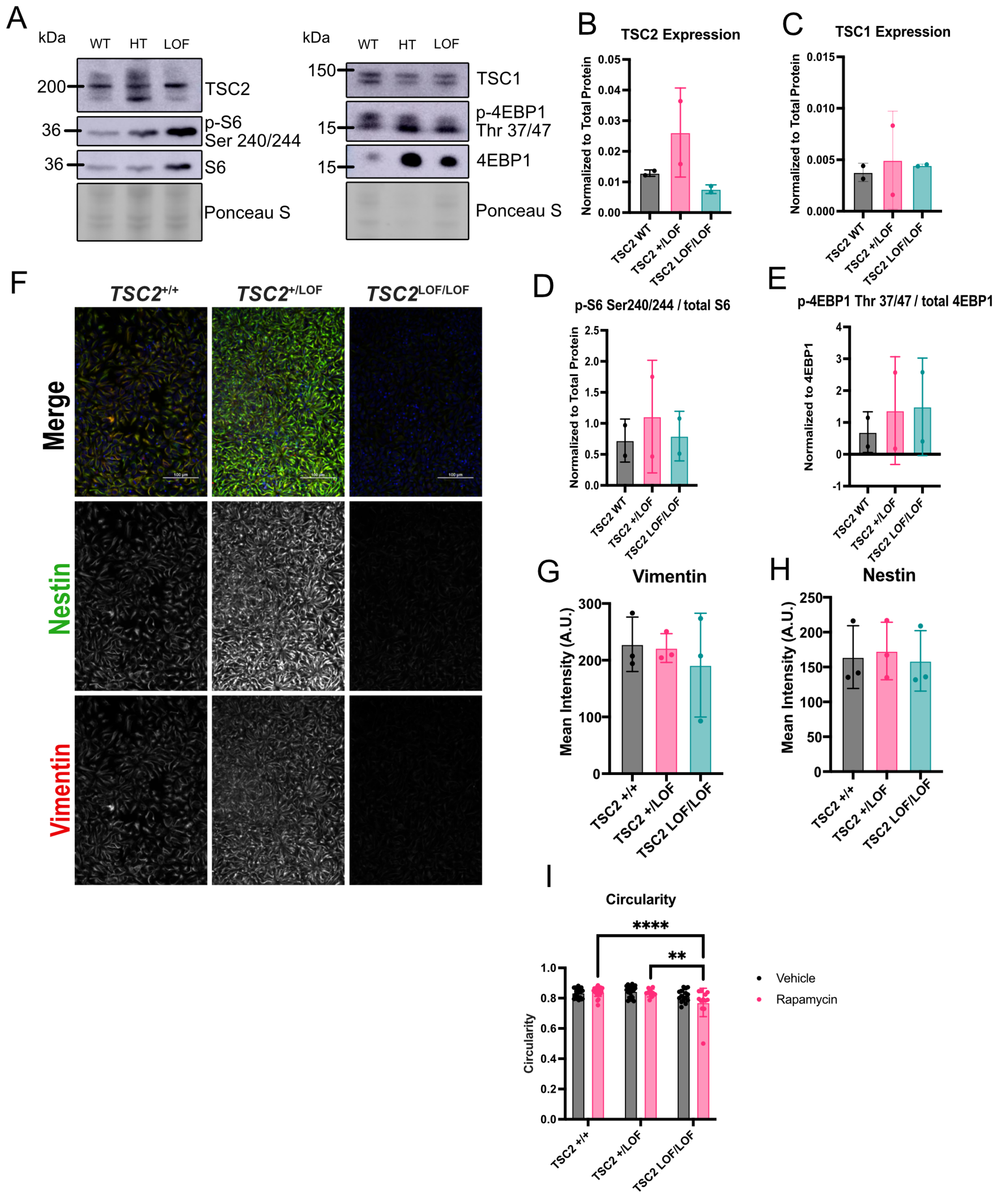


**Supplemental Figure 1: mTORC1 activity targets and neural progenitor markers**
(a) Representative immunoblot of day 10 monolayer culture showing expression of tuberin, hamartin, and downstream mTORC1 targets.

(b-e) Quantification of relative expression of tuberin, hamartin, p-S6 (ser 240/244), and p-4EBP-1 (thr 36/47) across multiple differentiations. Error bars = mean with SD.
(f) Representative immunofluorescence maximum intensity images of day 10 neural monolayer cultures showing expression of nestin (green), vimentin (red), and Hoechst (blue). Nestin and vimentin single channel greyscale images. Scale bars = 100 μm.
(g) Quantification of mean intensity (A.U.) of vimentin in day 10 monolayer culture. N = 3 independent differentiations. No significance [ANOVA]. Error bars = mean with SD.
(h) Quantification of mean intensity (A.U.) of nestin in day 10 monolayer culture. N = 3 independent differentiations. No significance [ANOVA]. Error bars = mean with SD.
(i) Quantification of circularity of lumens in neural rosettes in Fig 2.1f. Rapamycin +/+ vs LOF/LOF **** p = <0.0001, Rapamycin +/LOF vs LOF/LOF ** p = 0.0038, genotype **** p = <0.0001. [two way ANOVA]. Error bars = mean with SD.

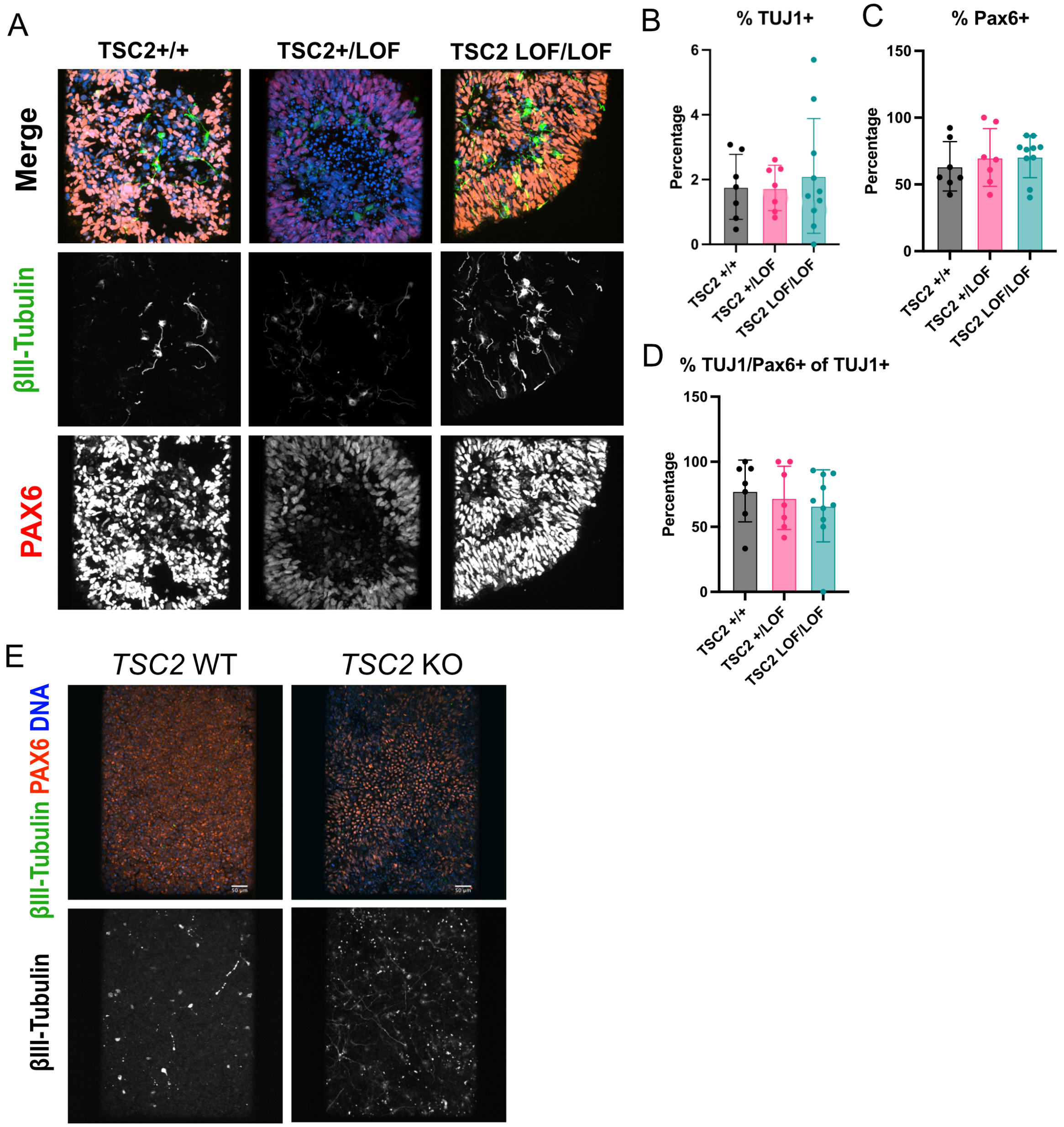


**Supplemental Figure 2: Neural Progenitor and Neuronal Markers in day 9 organoids**

(a) Representative immunofluorescence maximum intensity images of day 9 organoid neural cultures showing expression of β-III-tubulin (green), PAX6 (orange), and Hoechst (blue) to identify neural progenitor cells and neurons. β-III-tubulin and PAX6 single channel greyscale images. Scale bars = 100 μm.

(b) Quantification of percentage of β-III-tubulin positive soma. N = 9 independent organoids from multiple differentiations. No significance [ANOVA]. Error bars = mean with SD.

(c) Quantification of percentage of PAX6 positive nuclei. N = 9 independent organoids from multiple differentiations. No significance [ANOVA]. Error bars = mean with SD.

(d) Quantification of percentage of β-III-tubulin positive soma that are co-positive with PAX6. N = 9 independent organoids from multiple differentiations. No significance [ANOVA]. Error bars = mean with SD.
(e) Representative immunofluorescence maximum intensity images of day 9 organoid neural cultures showing expression of β-III-tubulin (green), PAX6 (orange), and Hoechst (blue) to identify neural progenitor cells and neurons. β-III-tubulin single channel greyscale images. Scale bars = 50 μm.

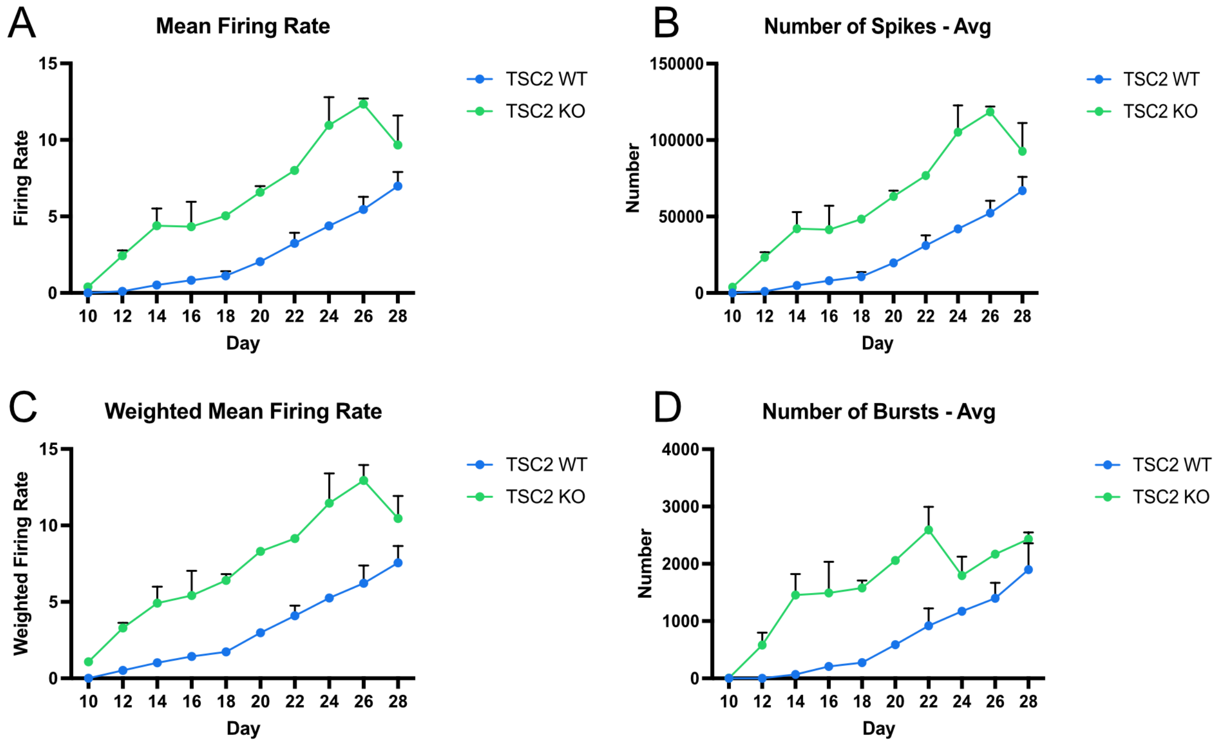


**Supplemental Figure 3: Increased electrical activity in TSC2 knock out cells between day 10 to day 28 of neural differentiation.**
(a) Quantification of mean firing rate over time. N = 16 wells over 2 independent differentiations. Error bars = mean with SD.
(b) Quantification of average number of spikes over time. N = 16 wells over 2 independent differentiations. Error bars = mean with SD.
(c) Quantification of weighted mean firing rate over time. N = 16 wells over 2 independent differentiations. Error bars = mean with SD.
(d) Quantification of average number of bursts over time. N = 16 wells over 2 independent differentiations. Error bars = mean with SD.

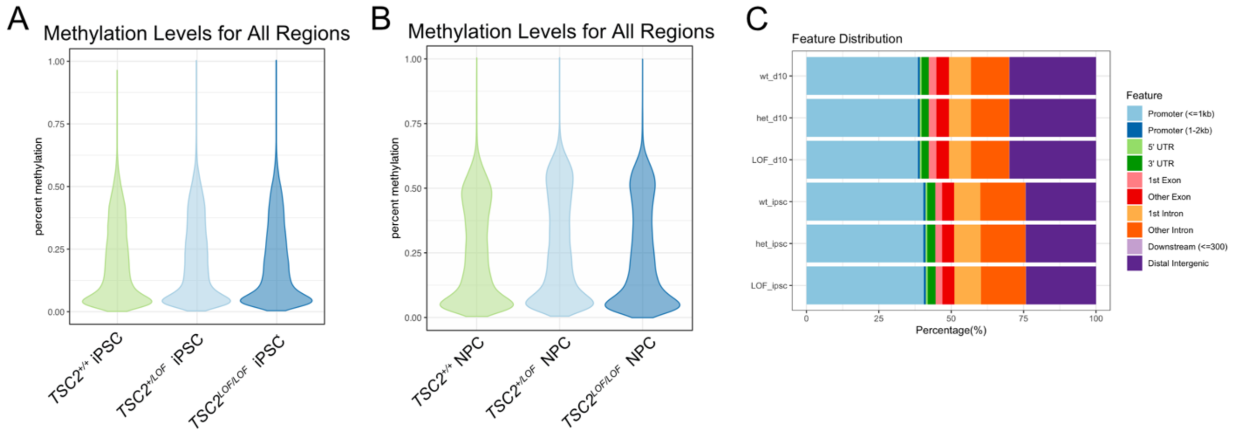


**Supplemental Figure 4: Global methylation levels in iPSCs and day 10 NPCs.**
(a) Violin plot showing global methylation levels for iPSCs for each genotype.
(b) Violin plot showing global methylation levels for day 10 NPCs for each genotype.

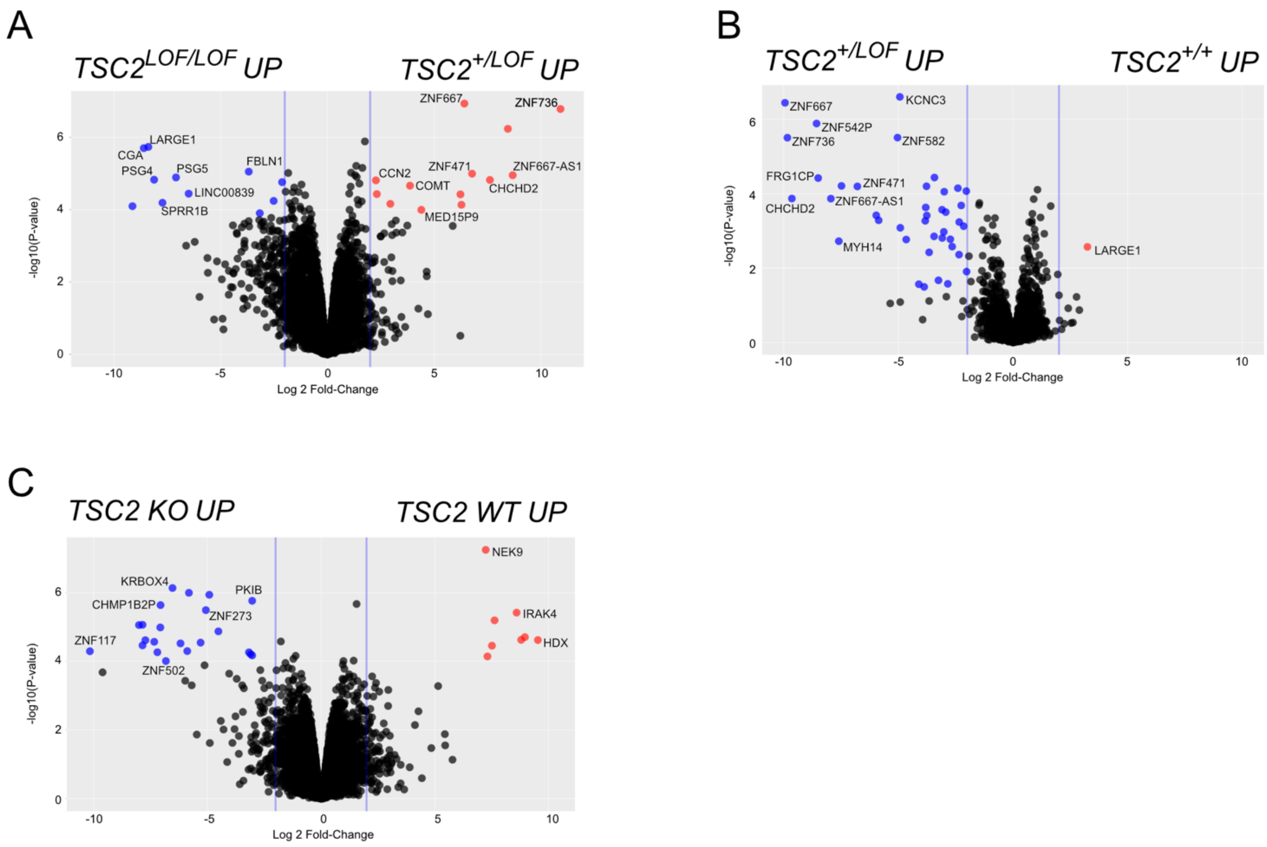


**Supplemental Figure 5: Differential gene expression in day 10 NPCs.**
(a) Volcano plot of differentially transcribed genes in *TSC2* +/LOF and *TSC2* LOF/LOF. Black dots are genes that are not both significant for fold change or adjusted p-value. Red dots are significantly upregulated in *TSC2* +/LOF by both fold change and adjusted p-value. Blue dots are significantly upregulated in *TSC2* LOF/LOF by both fold change and adjusted p-value.
(b) Volcano plot of differentially transcribed genes in *TSC2* +/+ and *TSC2* +/LOF. Black dots are genes that are not both significant for fold change or adjusted p-value. Red dots are significantly upregulated in *TSC2* +/+ by both fold change and adjusted p-value. Blue dots are significantly upregulated in *TSC2* +/LOF by both fold change and adjusted p-value.
(c) Volcano plot of differentially transcribed genes in *TSC2* WT and *TSC2* KO. Black dots are genes that are not both significant for fold change or adjusted p-value. Red dots are significantly upregulated in *TSC2* WT by both fold change and adjusted p-value. Blue dots are significantly upregulated in *TSC2* KO by both fold change and adjusted p-value.
